# MKK4 and MKK7 control degeneration of retinal ganglion cell somas and axons after glaucoma-relevant injury

**DOI:** 10.1101/2024.09.27.614559

**Authors:** Olivia J. Marola, Stephanie B. Syc-Mazurek, Sarah E. R. Yablonski, Peter G. Shrager, Simon W. John, Richard T. Libby

## Abstract

Glaucoma is characterized by programmed cell death of retinal ganglion cells (RGCs) after axonal injury. Several studies have shown the cell-intrinsic drivers of RGC degeneration act in a compartment-specific manor. Recently, the transcription factors JUN and DDIT3 were identified as critical hubs regulating RGC somal loss after mechanical axonal injury. It is possible somal DDIT3/JUN activity initiates axonal degeneration mechanisms in glaucoma. Alternatively, DDIT3/JUN may act downstream of inciting degenerative mechanisms and only drive RGC somal loss. The MAP2Ks MKK4 and MKK7 control all JNK/JUN activity and can indirectly activate DDIT3. Furthermore, MKK4/7 have been shown to drive RGC axonal degeneration after mechanical axonal injury. The present work investigated whether JUN and DDIT3, or their upstream activators MKK4 and MKK7, control degeneration of RGC axons and somas after glaucoma-relevant injury. *Ddit3*/*Jun* deletion did not prevent axonal degeneration in ocular hypertensive DBA/2J mice but prevented nearly all RGC somal loss. Despite robust somal survival, *Ddit3/Jun* deletion did not preserve RGC somal viability (as assessed by PERG decline and soma shrinkage) in DBA/2J mice or after glaucoma-relevant mechanical axonal injury. In contrast, *Mkk4/7* deletion significantly lessened degeneration of RGC somas and axons, and preserved somal function and size after axonal injury. In summary, activation of MKK4 and MKK7 appears to be the inciting mechanism governing death of the entire RGC after glaucoma-relevant injury; driving death of the RGC soma (likely through activation of DDIT3 and JUN), decline in somal viability, and axonal degeneration via DDIT3/JUN-independent mechanisms.

## Introduction

Vision loss in glaucoma is caused by progressive loss of retinal ganglion cells (RGCs). The most important risk factors for developing glaucoma are age and elevated intraocular pressure (IOP). After prolonged elevated IOP, RGC axons are injured at the optic nerve head—ultimately leading to programmed cell death. Much progress has been made in dissecting the mechanisms of RGC death after glaucoma-relevant injury. The pro-apoptotic molecule BAX was shown to be required for RGC somal death after chronic ocular hypertension (the DBA/2J mouse model of glaucoma). However, BAX was not required for axonal degeneration (1). Thus, RGC degeneration is compartmentalized after axonal injury; RGC somal and axonal degeneration are governed by distinct molecular mechanisms. Furthermore, the inciting mechanism triggered by ocular hypertension that leads to both axonal and somal degeneration must ultimately cause somal BAX activation, making upstream activators of BAX attractive targets of investigation.

Recently, the BAX-inducing transcription factors JUN and DNA-damage inducible transcript 3 (DDIT3, also known as CHOP) were identified as the critical regulators of RGC somal loss after glaucoma-relevant injury (2–4). *Ddit3* and *Jun* deletion provided near-complete protection to RGC somas after controlled optic nerve crush (CONC) (2), an acute model of mechanical axonal injury. Like *Bax* deletion, *Ddit3*/*Jun* deletion did not prevent RGC axonal degeneration after CONC (2), suggesting degenerative DDIT3/JUN activity is restricted to the RGC soma. Several lines of evidence have indicated axonal degeneration mechanisms are critical to initiating pathways driving both somal and axonal degeneration in glaucoma (5–8). However, it has also been suggested that degenerative mechanisms initiated in the soma are essential for driving axonal degeneration (9, 10). Somal JUN activity has been suggested to play a role in anterograde axonal injury signaling (9), and DDIT3 has been suggested to promote axonal degeneration, presumably as a result of its somal function (11). Individual deletions of *Ddit3* (4) or *Jun* (3) were not sufficient to prevent axonal degeneration in DBA/2J mice. However, it is possible DDIT3 and JUN play redundant and compensatory roles in initiating axonal degeneration. Given the near-complete somal protection conferred by combined *Jun*/*Ddit3* deletion after CONC (2), it is feasible somal DDIT3 and JUN act together to integrate somal and axonal degeneration cascades and thus govern death of the entire RGC in the context of ocular hypertension. Alternatively, it is also possible that DDIT3 and JUN act as critical downstream hubs regulating only RGC somal loss. If this is the case, the inciting mechanism governing degeneration of the entire RGC lies upstream of somal DDIT3 and JUN activation.

MAP2Ks 4 and 7 (MKK4/7) are known to act upstream of both DDIT3 and JUN activation. In fact, MKK4/7 are the only kinases known to activate the JUN-N-terminal kinases (JNKs 1, 2, and 3, also known as MAPKs 8, 9, and 10). Thus, MKK4/7 activation acts as a bottleneck for JNK and subsequent JUN activation (12, 13). As upstream effectors, MKK4/7 control several molecular mechanisms in addition to DDIT3/JUN. Through the JNKs, MKK4/7 can control JUN/DDIT3-independent cell death pathways, including NMNAT2 degradation (14, 15), SARM1 activation (16), and BH3-only protein activation (17). MKK4/7 also control JNK-independent degenerative pathways, including p38 (MAPK11-14) signaling (18). Individually, *Mkk4* or *Mkk7* deletion afforded significant but incomplete protection to RGC somas after CONC (42% and 17% protection, respectively) (19). Neither *Mkk4* nor *Mkk7* deletion alone was sufficient to prevent all JNK and JUN activation in RGC axons and somas (19), suggesting at least partial compensatory activity. Excitingly, blocking both MKK4 and MKK7 activity preserved morphological integrity of RGC axons after axonal injury (16). Therefore, MKK4/7 signaling may be a critical early mechanism governing RGC somal death via DDIT3/JUN activation and may also regulate axonal degeneration. Here, we assessed the importance of DDIT3 and JUN, along with their upstream activators MKK4 and MKK7, in RGC degeneration after glaucoma-relevant injury. Importantly, we investigated the role of these molecules in mitigating not only morphologically observable cell loss, but also deterioration of viability and gross function.

## Methods

### Mice

*Ddit3* null alleles (20) (Jackson Laboratory, Stock# 005530), floxed alleles of *Jun* (21) (*Jun^fl^*), and the Six3-cre transgene (22) (Jackson Laboratory, Stock# 019755) were backcrossed >10 times to both the C57BL/6J genetic background (>99% C57BL/6J) and the DBA/2J background (>99% DBA/2J). *Jun^fl^* alleles were recombined in the optic cup using Six3-cre. Floxed alleles of *Mkk4* (23) and *Mkk7* (24) were backcrossed >10 times to the C57BL/6J background and were recombined from retinal neurons via bilateral intravitreal delivery of AAV2.2-Cmv-cre-Gfp or AAV2.2-Cmv-Gfp with no cre (UNC vector core). Mice were fed chow and water ad libitum and were housed on a 12-hour light-to-dark cycle. All experiments were conducted in adherence with the Association for Research in Vision and Ophthalmology’s statement on the use of animals in ophthalmic and vision research and were approved by the University of Rochester’s University Committee on Animal Resources.

### Experimental rigor and statistical analysis

For all procedures, the experimenter was masked to genotype and condition. Roughly equal numbers of male and female mice were used for each experimental group. Animals were randomly assigned to experimental groups. Before experiments were performed, it was established that animals with pre-existing abnormal eye phenotypes (e.g. displaced pupil, cataracts) would be excluded from the study.

Data were analyzed using GraphPad Prism9 software. Power calculations were performed before experiments were conducted to determine appropriate sample size. The comparison of the percent of optic nerves at each grade between genotypes was analyzed using a Chi-square test. Data from experiments designed to test differences between two groups were subjected to an F test to compare variance and a Shapiro-Wilk test to test normality to ensure appropriate statistical tests were utilized. For normally distributed data with equal variance, a two-tailed independent samples *t* test was utilized. Data from experiments designed to test differences among more than two groups across one condition were subjected to a Brown-Forsythe test to compare variance and a Shapiro-Wilk test to test normality to ensure an appropriate statistical test was utilized. Normally distributed data with equal variance were analyzed using a one-way ANOVA followed by Holm-Sidak’s post-hoc test. Non-normally distributed data were analyzed using a Kruskal-Wallis test with Dunn’s post-hoc test. Data from experiments designed to detect differences among multiple groups and across two conditions were analyzed using a two-way ANOVA followed by Holm-Sidak’s post-hoc test. Data from experiments designed to test differences among multiple groups across more than two conditions were analyzed using a three-way ANOVA followed by Holm-Sidak’s post-hoc test. For these statistical tests, every possible comparison was made when relevant, and multiplicity adjusted *P* values are reported. In all cases, data met the assumptions of the statistical test used. *P* values <0.05 were considered statistically significant. Throughout the manuscript, results are reported as mean± standard error of the mean (SEM).

### Full field and pattern electroretinograms

Pattern and full-field electroretinography (PERGs and ERGs, respectively) was conducted using Diagnosys LLC’s Celeris rodent ERG system according to manufacturer’s instructions. Briefly, mice were dark-adapted for 60 minutes. Mice were anaesthetized with an intraperitoneal injection of 0.05 ml/10 g solution containing 20 mg/mL ketamine and 2 mg/mL xylazine. Hypromellose GenTeal (0.3%, Novartis Pharmaceuticals Corporation, NDC 0078-0429-47) was applied to the eyes before placement of electrodes. PERGs were obtained using 50cd/m^2^ mean luminance with spatial frequency 0.155 cycles/degree with 100% contrast. A total of 600 sweeps were recorded and averaged per eye. ERGs were obtained with 1 cd s/m^2^ luminance.

### Other surgeries and procedures

Controlled optic nerve crush (CONC) (1, 19, 25), intraocular pressure measurement (3, 25, 26), and intravitreal injections (27, 28) were performed as previously described. CONC was performed at least 28 days after intravitreal delivery of AAV2.2-Cmv-cre-gfp or AAV2.2-Cmv-gfp to allow for recombination and endogenous protein degradation.

### Compound action potentials

Compound action potentials were recorded as previously described (2, 29, 30). 5 days following CONC, animals were euthanized with CO_2_ asphyxiation and optic nerves were dissected free. Fresh optic nerves were transferred to a chamber of artificial cerebral spinal fluid (ACSF) aerated with 95% O_2_/5%CO_2_ for at least 60 minutes before recording. Optic nerves were transferred to a temperature-controlled chamber perfused with ACSF bubbled with 95% O_2_ / 5% CO_2_. Nerves were drawn into glass pipet suction electrodes (filled with ACSF) at each end for stimulation and recording. The recording pipet resistance was measured before (17-20 KΩ) and after (29-34 KΩ) insertion of the nerve and monitored continuously during the experiment. The ratio of this resistance during each sweep divided by the resistance of the pipet alone allowed a normalization of the amplitude of the compound action potential (CAP) to our standard ratio of 1.7 (30, 31). This corrects for any drift in the seal resistance during an experiment. Signals were fed to one input of an AC differential amplifier of our design. The second input came from a pipet electrode placed near the recording electrode. This served to subtract much of the stimulus artifact. Stimuli of 50μs duration were delivered by an optically isolated constant current unit (WPI, Sarasota FL) driven by the computer. Stimulus currents were monitored by a linear optically coupled amplifier of our design. All signals were electronically low pass filtered with a cutoff of 10 kHz. All records were taken at 37 ± 0.5 °C.

### Histology, nerve grading, and immunofluorescence

Optic nerve processing for plastic sectioning and optic nerve severity grading (3, 4) were performed as previously described. Immunofluorescence of whole-mounted retinas (4, 19, 25) and cryosectioned optic nerves (29, 30) were performed as previously described using the following antibodies: rabbit anti-cCASP3 (AF835, R&D, 1:1000), rabbit anti-RBPMS (GTX118619, GeneTex, 1:250), chicken anti-GFP (ab13970-100, Abcam, 1:1000), rabbit anti-pJNK (4668S , Cell Signaling, 1:250), rabbit anti-pJUN (3270S, Cell Signaling, 1:250), donkey anti-rabbit (A31572 and A-21206, ThermoFisher, 1:1000), donkey anti-mouse (A31570, ThermoFisher, 1:1000), and donkey anti-chicken (703-545-155, Jackson ImmunoResearch, 1:1000). RGC soma sizes (measured using images assessed for RGC soma survival) were quantified using ImageJ by using a Gaussian blur filter with a sigma of 4, converting images to binary with an automatic triangle thresholding setting allowing detection of RGC somas. Somas were separated using the watershed function. Somas were defined as an area ≥30µm^2^ and ≥.2 circularity. RGCs cut off at the boarder of the image were excluded from analysis. Average soma area per image was measured, and 8 images were averaged per retina.

## Results

### Ddit3 and Jun control RGC somal loss after ocular hypertension

DDIT3 and JUN together have been shown to regulate death of RGC somas after CONC, potentially additively (2). It has yet to be determined whether somal DDIT3 and JUN activity initiates axonal degeneration mechanisms. To determine whether both JUN and DDIT3 act in tandem to elicit RGC axonal degeneration in the context of glaucoma, *Ddit3* and/or *Jun* were deleted from the full body and neural retina, respectively, from ocular hypertensive DBA/2J mice.

Neurodegeneration in the DBA/2J model of glaucoma is dependent upon elevated IOP (32–37). Some genetic manipulations have been reported to lower IOP in DBA/2J mice (25). However, with Six3-cre, *Jun* is not deleted in the structures in the anterior segment of the eye that control IOP, and D2.Six3-cre^+^*Jun^fl/fl^*mice did not have altered IOP compared to WT controls (3). Furthermore, full-body deletion of *Ddit3* did not alter the IOP profile of the DBA/2J mouse model (4). To ensure combined *Ddit3/Jun* deletion did not lessen IOP elevation, IOPs were measured from D2.*Ddit3^+/?^Jun^+/?^*(WT) and D2.Six3-cre^+^*Ddit3^-/-^Jun^fl/fl^* (*Ddit3/Jun^-/-^*) mice at 5M, 9M, 10.5M, and 12M of age. Mice of both genotypes had elevated IOP at each timepoint compared to 5M, and IOP was not lowered compared to WT at each timepoint measured. Therefore, *Ddit3*/*Jun* deletion did not substantially alter the profile of ocular hypertension typical of the DBA/2J model (Fig. 1A).

**Fig. 1.**
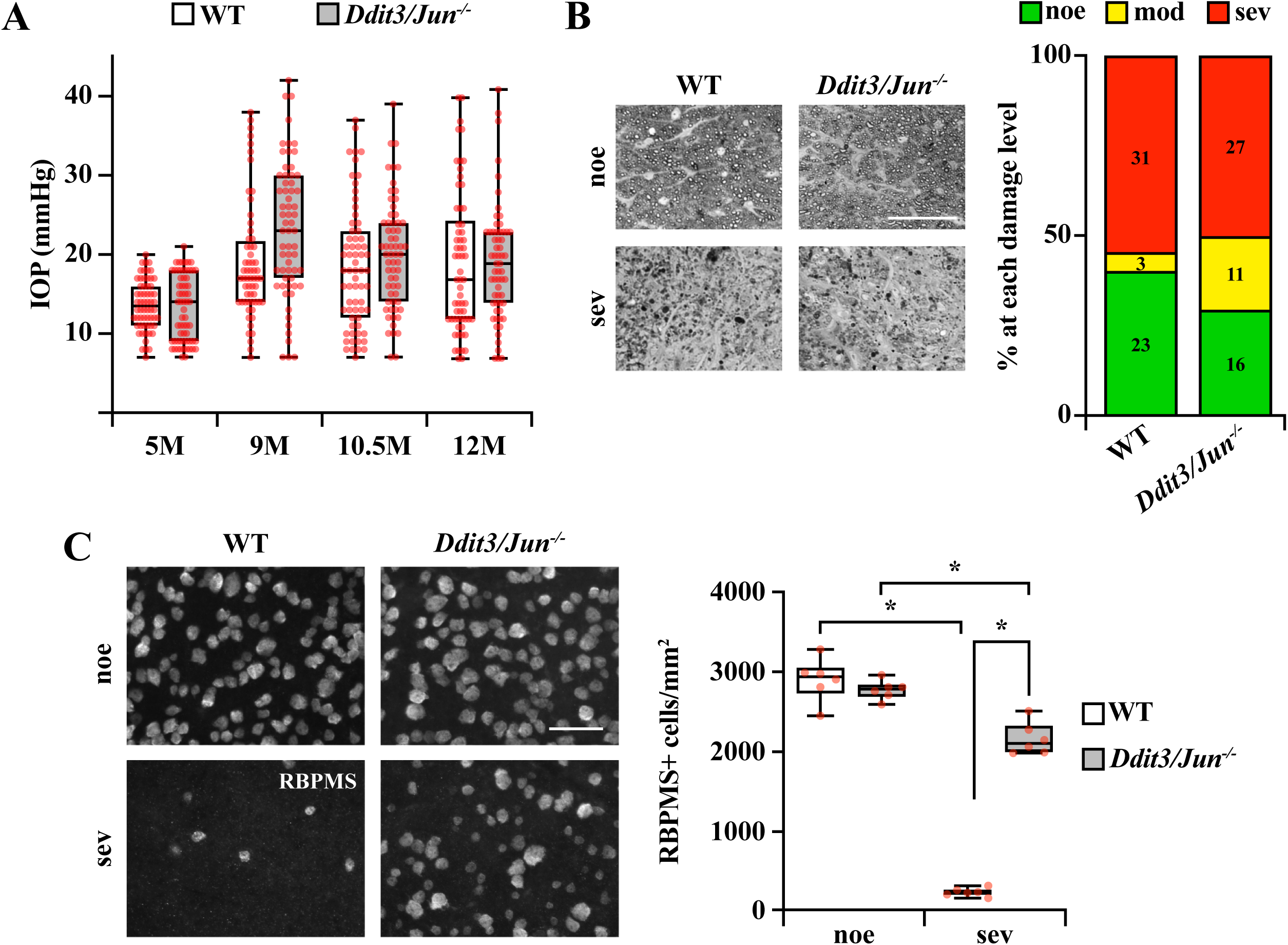
| *Ddit3/Jun* deletion did not prevent axonal degeneration but robustly improved RGC somal survival in DBA/2J mice. **A.** Longitudinal intraocular pressure (IOP) measurements from WT and *Ddit3/Jun^-/-^*eyes at 5, 9, 10.5, and 12 months of age. Both genotype groups had significantly elevated IOPs at 9M, 10.5M, and 12M compared to 5M (*P<*0.001, *n≥*60 for each comparison). At these timepoints, *Ddit3/Jun^-/-^* eyes did not have a statistically significant reduction in IOP compared to WT, although *Ddit3/Jun^-/-^* eyes had slightly higher IOPs at 9M of age compared to WT (*n≥*60, \**P=*0.004, Two-way ANOVA, Holm-Sidak’s *post hoc*). **B.** Examples of optic nerve cross sections with no or early (noe) and severe (sev) glaucomatous damage from WT and *Ddit3/Jun^-/-^*mice and percentages of optic nerves with noe, moderate (mod) and sev glaucomatous damage in 12M WT and *Ddit3/Jun^-/-^* mice. *Ddit3/Jun* deletion did not afford protection to RGC axons and in fact slightly worsened axonal degeneration (*n≥*54, *P=*0.049, Chi square test). **C.** Representative retinal flat mounts immunoassayed for RBPMS and quantification of RBPMS+ cells from 12M WT and *Ddit3/Jun^-/-^* retinas with corresponding noe or sev optic nerves. Sev *Ddit3/Jun^-/-^* retinas had 77.0±3.1% improved RGC survival compared to WT controls. RBPMS+ cells/mm^2^±SEM for WT and *Ddit3/Jun^-/-^* respectively: Noe: 2899.6±111.0, 2765.4±49.8; Sev: 226.7±20.8, 2151.5±83.0 (*n=*6, \**P*<0.001, Two-way ANOVA, Holm-Sidak *post-hoc*). Scale bars, 50μm.

To determine whether DDIT3 and JUN control death of the entire RGC after ocular hypertension, *Ddit3/Jun^-/-^*and WT control optic nerves were assessed for axonal degeneration at 12M—a timepoint at which roughly 50% of DBA/2J optic nerves will have severe optic nerve damage (26). *Ddit3/Jun* deletion did not lessen instances of severe glaucomatous neurodegeneration. In fact, *Ddit3/Jun^-/-^* mice had slightly worse outcomes relative to WT controls (Fig. 1B). Therefore, somal DDIT3 and JUN did not act in tandem to perpetuate axonal degeneration after chronic ocular hypertension.

Several molecules contribute to degeneration of the soma, but not the axon, after glaucoma-relevant injury (1, 3, 4). To determine whether DDIT3 and JUN play an important role in RGC somal degeneration after severe axonal injury, 12M *Ddit3/Jun^-/-^* and WT retinas with corresponding severe optic nerves were assessed for RGC somal survival. *Ddit3/Jun* deletion conferred robust (77%) protection to RGC somas in retinas with severe optic nerve degeneration (Fig. 1C). These data suggest DDIT3 and JUN are the critical regulators of somal death but do not contribute to axonal degeneration in glaucoma.

### Despite somal survival, Ddit3/Jun deletion did not prevent PERG amplitude decline or somal shrinkage

Ocular hypertension is known to cause impaired RGC somal gross potentials (5–7, 38, 39), even before the onset of detectable axonal damage (39, 40). Others have shown neuroprotective treatment or genetic manipulation also preserved physiological activity in glaucoma-relevant models (5–7, 38). Given *Ddit3*/*Jun* deletion protected most RGC somas after chronic ocular hypertension (Fig. 1C), it remained important to determine whether surviving *Ddit3/Jun^-/-^*RGC somas retained physiological function. To test this, pattern electroretinograms (PERGs) were longitudinally recorded from WT, D2.*Ddit3^-/-^Jun^+/?^* (*Ddit3^-/-^*), D2.Six3-cre^+^ *Ddit3^+/?^Jun^fl/fl^* (*Jun^-/-^*), and *Ddit3/Jun^-/-^* animals at 5M (before the onset of IOP elevation and RGC degeneration (26)), 9M (when IOP is elevated, but morphologically detectable RGC degeneration has not yet occurred (26)), and 12M of age (when roughly 50% of eyes have severe glaucomatous neurodegeneration (26), Fig. 1B). D2.*Gpnmb^+^* (*Gpnmb^+^*) animals were assessed as an age- and background-matched normotensive control (35).

Compared to non-glaucomatous *Gpnmb^+^* age-matched controls, PERG amplitude significantly declined in all genotype groups over time (Fig. 2A, B). As previously observed, PERG amplitude significantly declined by 9M for DBA/2J mice compared to *Gpnmb^+^* age-matched controls (39), and *Ddit3/Jun* deficiency did not prevent this decline. Similar results were observed at 12M. Thus, despite conferring robust protection to RGC somas, *Ddit3*/*Jun* deletion did not prevent loss of RGC somal function. Notably, PERG amplitude decline did not appear to be caused by photoreceptor or bipolar cell dysfunction or improper light penetration. ERG a- and b-wave amplitudes declined slightly with age for all genotype groups, including *Gpnmb^+^* eyes, but not nearly to the same extent as PERG amplitude decline (Fig. 2C, D). %PERG and ERG a- and b-wave amplitudes relative to respective 5M controls are listed in Table 1.

**Fig. 2.**
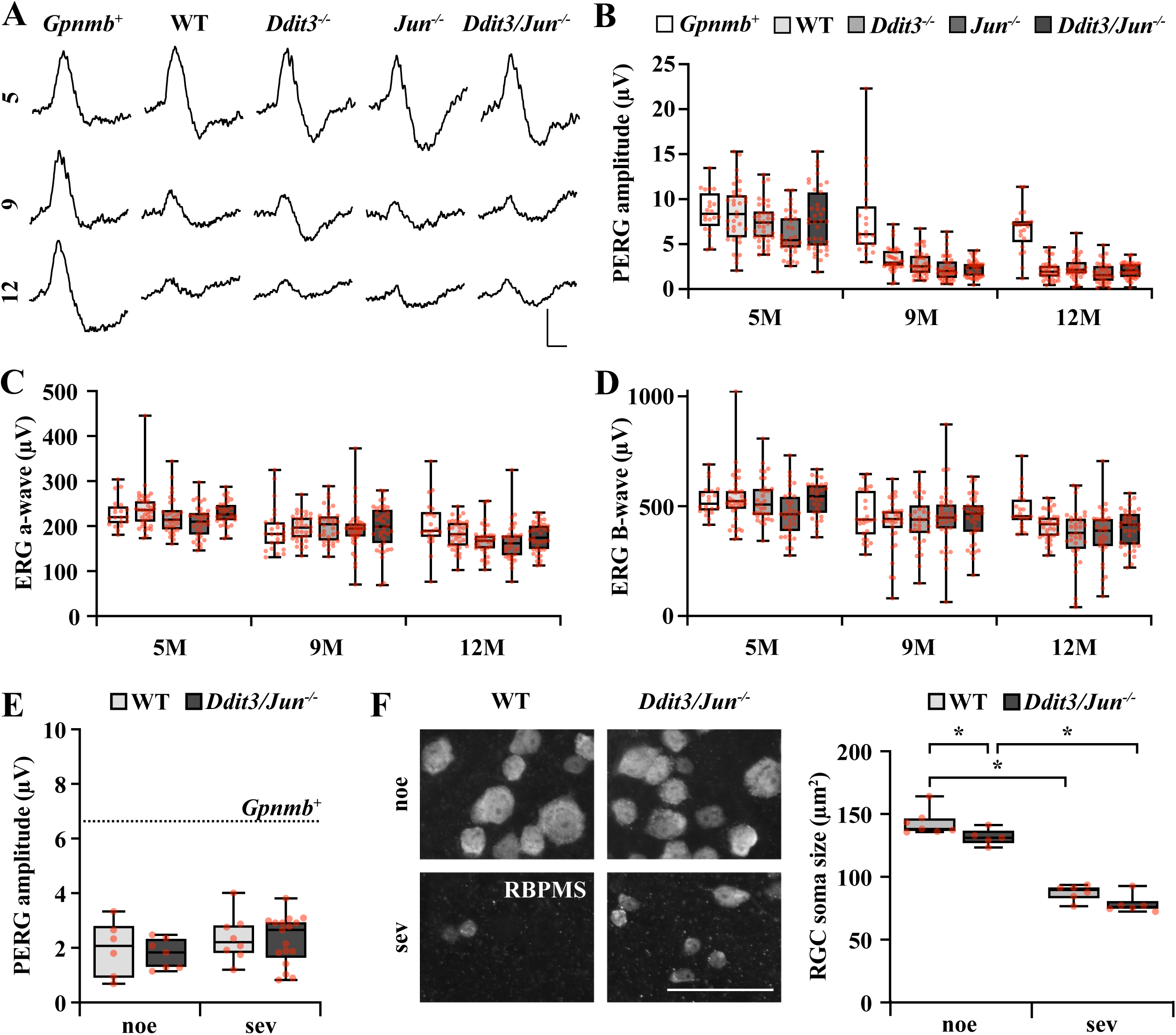
| *Ddit3/Jun* deletion did not prevent decline of PERG amplitudes in DBA/2J mice. **A.** Representative pattern electroretinography (PERG) traces and amplitude quantification (**B**) from Gpnmb^+^, WT, Ddit3^-/-^, Jun^-/-^, and Ddit3/Jun^-/-^ mice at 5, 9, and 12M. Normotensive Gpnmb^+^ mice did not have significant decline in PERG amplitude at 9M (n≥21, P>0.05), but had a slight but significant decline in PERG amplitude by 12M compared to 5M (n≥21, P=0.007), Ocular hypertensive mice of all genotype groups had significant PERG amplitude decline at 9M and 12M compared to 5M (n≥34, P<0.001, two-way ANOVA, Holm-Sidak’s post hoc). At 9M and 12M, each ocular hypertensive group’s PERG amplitude was significantly lower than normotensive Gpnmb^+^ controls (n≥20, P<0.001). No increase in PERG amplitude was observed between WT and Ddit3^-/-^, Jun^-/-^, or Ddit3/Jun^-/-^ groups at any timepoint measured. Scale bar: Y: 5μV, X: 100ms. **C.** Quantification of full-field ERG a-wave and b-wave (**D**) amplitudes. By 12M, each ocular hypertensive group had a slight but significant decline of electroretinography (ERG) a and b-wave amplitudes compared to 5M (n≥34, P<0.001, two-way ANOVA, Holm-Sidak’s post hoc), but not nearly to the same extent as PERG amplitude decline (**A**). Percentage of PERG and ERG amplitude declines at 9 and 12M are listed for each group in Table 1. **E.** PERG amplitude quantifications from 12M WT and *Ddit3/Jun^-/-^* retinas with noe and sev optic nerve damage. Neither genotype nor optic nerve damage level influenced PERG amplitude (n≥6, *P*>0.05, two-way ANOVA). Of note, PERG amplitudes were significantly reduced compared with 12M *Gpnmb*^+^ (*n*=21, 7.2±0.6) regardless of genotype or level of axonal damage. PERG amplitude (μV)±SEM: WT noe: 2.0±0.4, *Ddit3/Jun^-/-^* noe: 1.8±0.2, WT sev: 2.3±0.3, *Ddit3/Jun^-/-^*sev: 1.9±0.2 (*n≥*6, *P<*0.001, one-way ANOVA, Holm-Sidak’s *post hoc*). Scale bar: Y: 5μV, X: 100ms. **F.** High-resolution images of retinal flat mounts immunoassayed for RBPMS (scale bar, 50μm) and quantification of average RGC soma size from WT and *Ddit3/Jun^-/-^* retinas with noe and sev glaucomatous damage. Both genotype groups had significant reductions in RGC soma size in sev glaucoma compared to respective noe controls (\**P*<0.001). While *Ddit3/Jun*^-/-^ noe retinas had slightly smaller RGCs (\**P*=0.044), *Ddit3/Jun* deletion did not attenuate RGC soma shrinkage in sev retinas. Soma size (μm^2^)±SEM from WT and *Ddit3/Jun^-/-^* retinas, respectively: noe: 143.1±3.9, 131.7±2.9; sev: 87.6±2.5, 78.3±3.0 (n≥5, two-way ANOVA, Holm-Sidak’s *post hoc*).

**Table 1.** | Amplitudes of PERGs, ERG a-waves, and ERG b-waves expressed as a percentage of respective 5M amplitudes

|  |  | *Gpnmb*<sup>+</sup> | WT | *Ddit3*<sup>-/-</sup> | *Jun*<sup>-/-</sup> | *Ddit3/Jun*<sup>-/-</sup> |
| --- | --- | --- | --- | --- | --- | --- |
| %PERG amplitude | 9M | 89.0 ± 9.9 | **39.8 ± 2.9\*** | **38.8 ± 3.1\*** | **38.3 ± 3.6\*** | **29.6 ± 2.1\*** |
|  | 12M | **77.2 ± 6.3** | **25.0 ± 2.1\*** | **31.7 ± 2.8\*** | **29.7 ± 3.1\*** | **27.4 ± 1.9\*** |
| %ERG a wave amplitude | 9M | **84.0 ± 4.2** | **82.5 ± 2.2** | 90.4 ± 2.9 | 92.0 ± 3.8 | **86.0 ± 3.5** |
|  | 12M | 89.0 ± 5.3 | **76.4 ± 2.2** | **75.5 ± 2.6\*** | **78.6 ± 3.6\*** | **76.4 ± 2.4\*** |
| %ERG b wave amplitude | 9M | **86.0 ± 3.9** | **79.7 ± 3.7** | **81.9 ± 3.6** | 95.2 ± 4.7 | **83.8 ± 3.2** |
|  | 12M | 91.5 ± 3.6 | **76.8 ± 2.0\*** | **68.3 ± 4.1\*** | **79.2 ± 4.6\*** | **75.0 ± 2.7\*** |

**Bold text:** $P < 0.05$ compared to respective 5M
\*: $P < 0.05$ compared to age-matched *Gpnmb*<sup>+</sup>

Notably, PERG amplitude decline did not depend on morphologically observed RGC loss. Regardless of genotype, 12M eyes with no or early and severe glaucomatous damage had similarly reduced PERG amplitudes relative to age-matched *Gpnmb^+^* controls (Fig. 2E). Interestingly, *Ddit3/Jun* deletion did not prevent ocular hypertension-induced shrinkage of the RGC soma (Fig. 2F), suggesting surviving RGCs are likely injured and/or undergoing metabolic stress (41, 42). Thus, despite conferring protection from somal loss, *Ddit3/Jun* deletion did not preserve RGC somal viability, at least as measured by gross potentials and soma size. These data suggest the mechanism(s) driving somal shrinkage and loss of gross potentials must act either upstream or independently of somal DDIT3/JUN activation.

### Mkk4 and Mkk7 deletion prevented somal and axonal degeneration after axonal injury

MKK4 and MKK7 (MAP2Ks 4 and 7) are known to control DDIT3 signaling and all JUN activation after injury, and thus may drive RGC somal death after axonal injury. In addition, MKK4/7 activate a variety of downstream targets independently of DDIT3/JUN activation, which may contribute to decline of somal viability and axonal degeneration. In fact, recent work has suggested the importance of both MKK4 and MKK7 in driving axonal degeneration (16, 43). Therefore, MKK4/7 activation may drive degeneration of the entire RGC. To test this possibility, *Mkk4^fl^* and *Mkk7^fl^* alleles were recombined from RGCs using intravitreal AAV2.2-delivered Cmv-cre (AAV2.2-Cmv-cre-Gfp) to generate animals with *Mkk4/7^-/-^* RGCs. AAV2.2 with no cre vector (AAV2.2-Cmv-Gfp) was intravitreally injected into *Mkk4^?^Mkk7^?^* mice to generate *Mkk4/7*^+/+^ controls.

To ensure sufficient recombination of *Mkk4/7* floxed alleles, activation of MKK4/7’s downstream targets was evaluated after CONC. Robust JNK activation in the optic nerve head and JUN activation in RGC somas occurs early after CONC (2, 44). Compared to *Mkk4/7^+/+^*controls, *Mkk4/7^-/-^* eyes had little appreciable JNK activation in the optic nerve head (Fig. 3A) and had a 91% reduction of RGCs with JUN activation (Fig. 3B) after CONC. Therefore, AAV2.2-delivered Cmv-cre effectively recombined *Mkk4/7* floxed alleles in RGCs. To determine whether MKK4 and MKK7 together play an important role in RGC somal loss after glaucoma-relevant injury, RGC soma survival was assessed for *Mkk4/7*^+/+^ and *Mkk4/7*^-/-^ mice after CONC. *Mkk4/7* deletion provided robust and sustained long-term protection to RGC somas after CONC—*Mkk4/7* deletion prevented the vast majority of caspase 3 activation 5 days post-CONC (Fig. 4A) and preserved ∼90% of RGC somas at both 14 days and 2 months post-CONC (Fig. 4B). Therefore, MKK4 and MKK7 controlled RGC somal death after glaucoma-relevant injury.

**Fig. 3.**
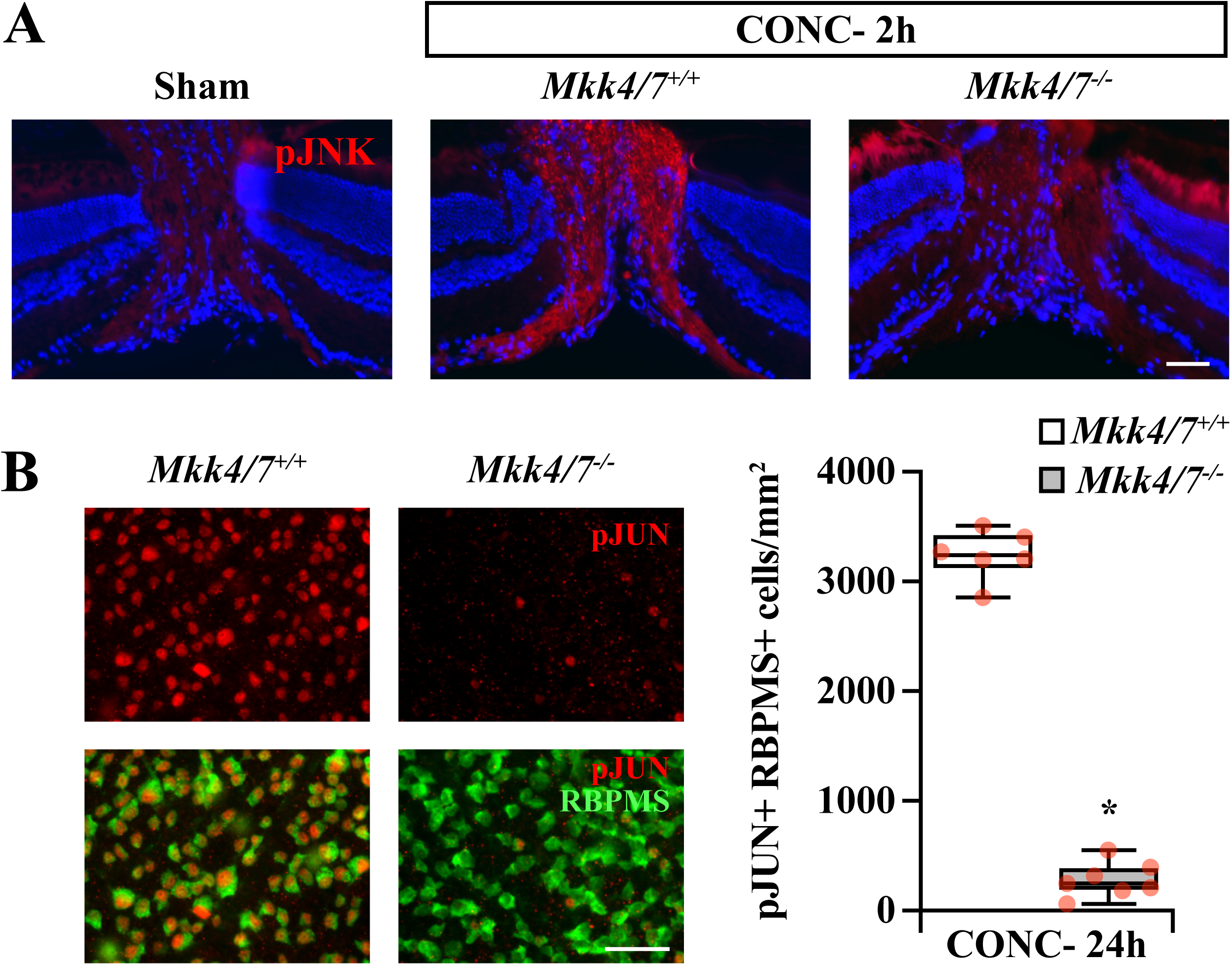
| Efficient RGC recombination of *Mkk4* and *Mkk7* floxed alleles with AAV2.2 delivered Cmv-cre. **A.** Representative sections of *Mkk4/7^+/+^* and *Mkk4/7^-/-^*optic nerve heads 2 hours after CONC immunoassayed for activated JNK (pJNK). JNK was markedly activated in *Mkk4/7^+/+^* optic nerves, but not in *Mkk4/7^-/-^* optic nerves, 2 hours after CONC (*n=*3). **B.** Representative *Mkk4/7^+/+^* and *Mkk4/7^-/-^*retinal flat mounts 24 hours after CONC immunoassayed for RBPMS (green) and activated JUN (pJUN, red) and quantification of pJUN+ RBPMS+ cells. RGC JUN activation was reduced by 91.4±1.8% in *Mkk4/7^-/-^*retinas. pJUN+RBPMS+ cells/mm^2^: *Mkk4/^+/+^*: 3239.6±91.9, *Mkk4/7^-/-^*: 278.0±59.8 (*n≥*6, \**P<*0.001, two-tailed *t* test). Scale bar, 50μm.

**Fig. 4.**
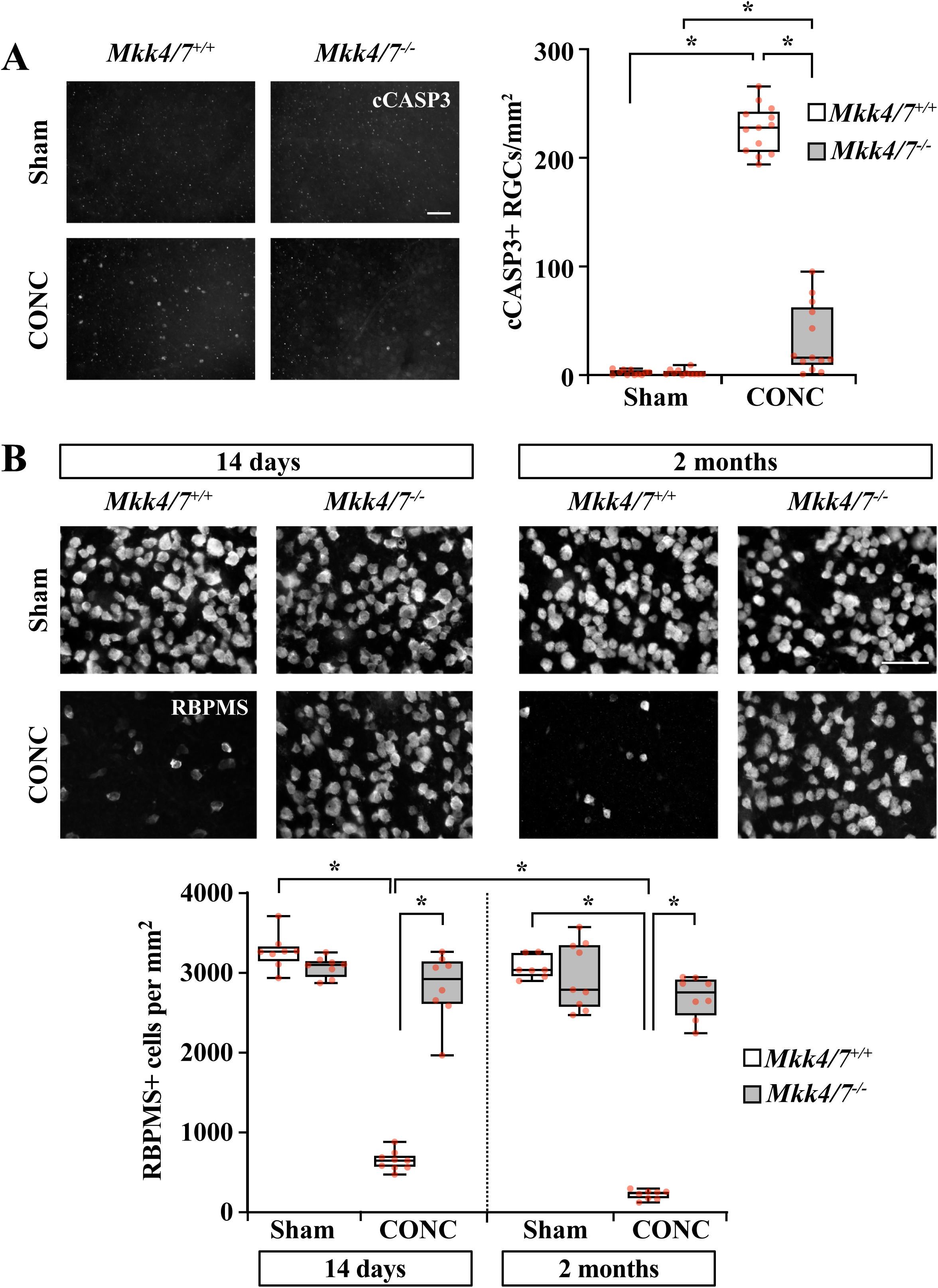
| *Mkk4/7* deletion protected nearly all RGC somas after axonal injury. **A.** Representative *Mkk4/7^+/+^* and *Mkk4/7^-/-^* retinal flat mounts 5 days after CONC immunoassayed for activated caspase 3 (cCASP3). *Mkk4/7* deletion prevented 85.6±3.9% of caspase 3 activation. cCASP3+RGCs/mm^2^±SEM for *Mkk4/7^+/+^*and *Mkk4/7^-/-^* respectively: Sham: 2.2±0.7, 2.4±0.8; CONC: 226.5±6.1, 32.5±8.8 (*n≥*11, \**P<*0.001, two-way ANOVA, Holm-Sidak’s *post hoc*). **B.** Representative *Mkk4/7*^+/+^ and *Mkk4/7*^-/-^ retinal flat mounts 14 days and 2 months after CONC immunoassayed for RBPMS and quantification of RBPMS+ cell survival. *Mkk4/7* deletion prevented 89.9±6.1% and 90.2±3.3% of RGC death at 14 days and 2 months, respectively. RBPMS+ cells/mm^2^±SEM for 14 days—*Mkk4/7^+/+^*Sham: 3264.7±77.8, *Mkk4/7^-/-^* Sham: 3067.8±45.1, *Mkk4/7^+/+^* CONC: 667.1±38.0, *Mkk4/7^-/-^* CONC: 2819.0±149.6. For 2 months—*Mkk4/7^+/+^* Sham: 3063.0±53.2, *Mkk4/7^-/-^* Sham: 3015.6±144.6, *Mkk4/7^+/+^* CONC: 222.6±21.0, *Mkk4/7^-/-^* CONC: 2690.7±91.7 (*n*≥7, *Mkk4/7^+/+^* CONC 14 days vs 2 months, \**P*<0.05; *Mkk4/7^+/+^*Sham vs CONC at 14 days and 2 months, \**P<*0.001; *Mkk4/7^+/+^* vs *Mkk4/7^-/-^* CONC at 14 days and 2 months: \**P*<0.001. Three-way ANOVA, Holm-Sidak’s *post hoc*). Scale bars, 50μm.

Recent evidence has suggested a role for MKK4/7 in driving Wallerian degeneration cascades after glaucoma-relevant injury (16, 43). To clarify the role of MKK4/7 in axonal degeneration, RGC axonal integrity was evaluated for *Mkk4/7^+/+^* and *Mkk4/7*^-/-^ optic nerves after glaucoma-relevant injury. Consistent with previous reports (16, 43), *Mkk4/7^-/-^* RGC axons had substantially fewer histological indications of degeneration (Fig. 5A). Importantly, *Mkk4/7* deletion preserved axonal physiological function as assessed by compound action potentials (CAPs, Fig. 5B, C). These data show that together, MKK4 and MKK7 control degeneration of both the RGC soma and axon after axonal injury.

**Fig. 5.**
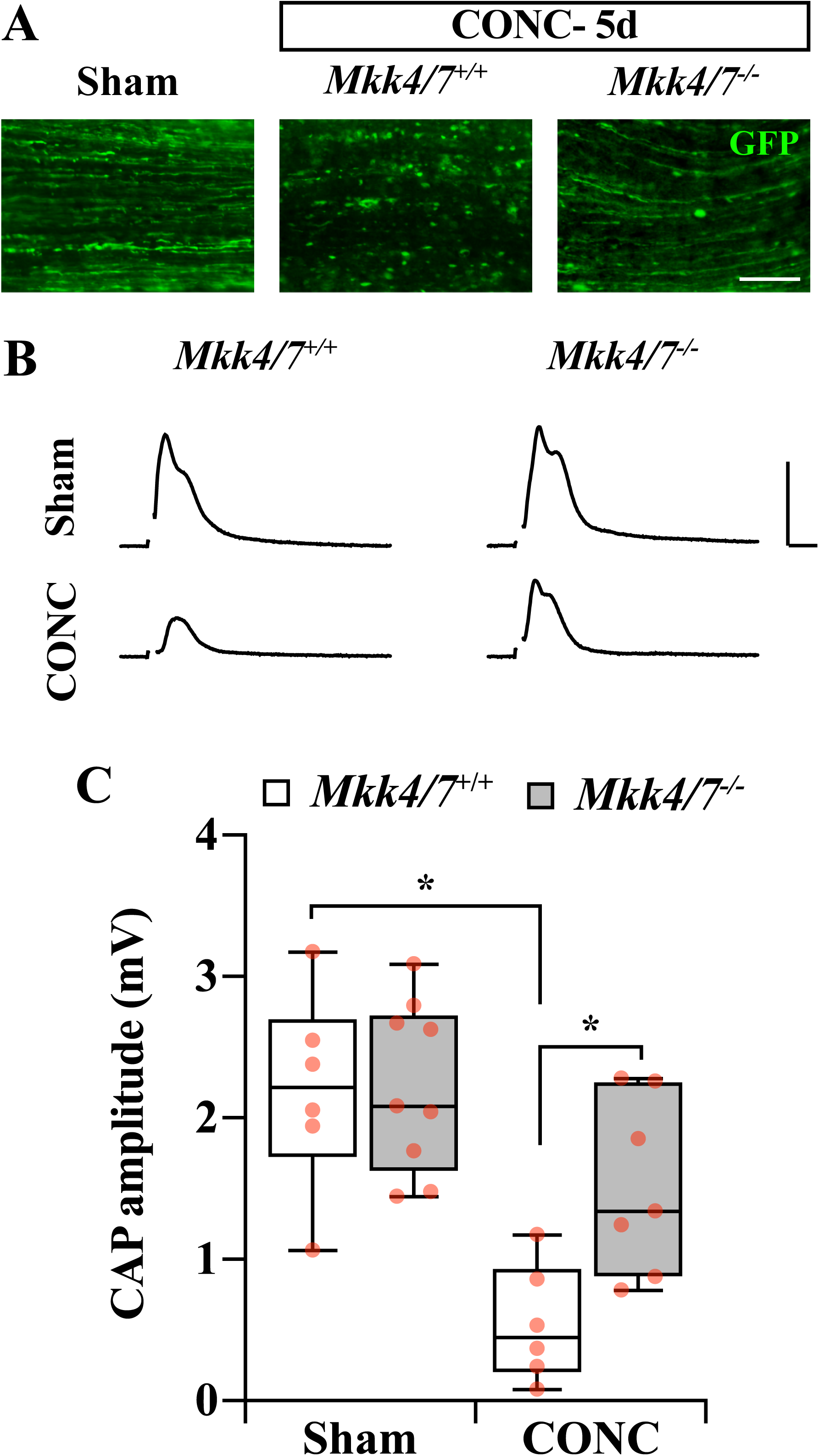
| *Mkk4/7* deletion prevented RGC axonal degeneration after axonal injury. **A.** Representative longitudinal sections of *Mkk4/7^+/+^* and *Mkk4/7^-/-^* optic nerves 5 days after CONC or Sham procedures. *Mkk4/7^-/-^* optic nerves had significantly improved axonal integrity compared to *Mkk4/7^+/+^* controls. Note: AAV2.2-Cmv-cre-Gfp and AAV2.2-Cmv-Gfp controls both induce GFP expression, therefore RGC axons were identified with GFP. Scale bar, 50μm. **B.** Representative compound action potential (CAP) traces from of *Mkk4/7^+/+^* and *Mkk4/7^-/-^*optic nerves 5 days after CONC or Sham procedures. Scale bar: Y: 2mV, X: 1ms. **C.** Quantification of CAP amplitudes from of *Mkk4/7^+/+^* and *Mkk4/7^-/-^* optic nerves 5 days after CONC or Sham. *Mkk4/7* deletion provided significant protection from CAP amplitude decline after CONC. CAP amplitude (mV)±SEM from *Mkk4/7^+/+^* and *Mkk4/7^-/-^,* respectively: Sham: 2.8±0.3, 2.8±0.2; CONC: 0.9±0.2, 2.0±0.3. (*n≥*6, \**P<*0.05, two-way ANOVA, Holm-Sidak’s *post hoc*).

Given PERG amplitude decline and soma shrinkage were not prevented with *Ddit3/Jun* deletion despite robust somal survival, it remained important to assess RGC somal and function size in *Mkk4/7^-/-^*eyes after glaucoma relevant injury. In striking contrast to *Jun/Ddit3* deletion, *Mkk4*/*7* deletion significantly attenuated PERG amplitude decline (Fig. 6A) and soma shrinkage (Fig. 6B) after CONC. These data suggest MKK4/7 govern not only somal and axonal survival, but also drive loss of viability and gross function. Thus, activation of MKK4/7 is likely a critical inciting event integrating mechanisms controlling death and degeneration of somal and axonal RGC compartments in the context of axonal injury.

**Fig. 6.**
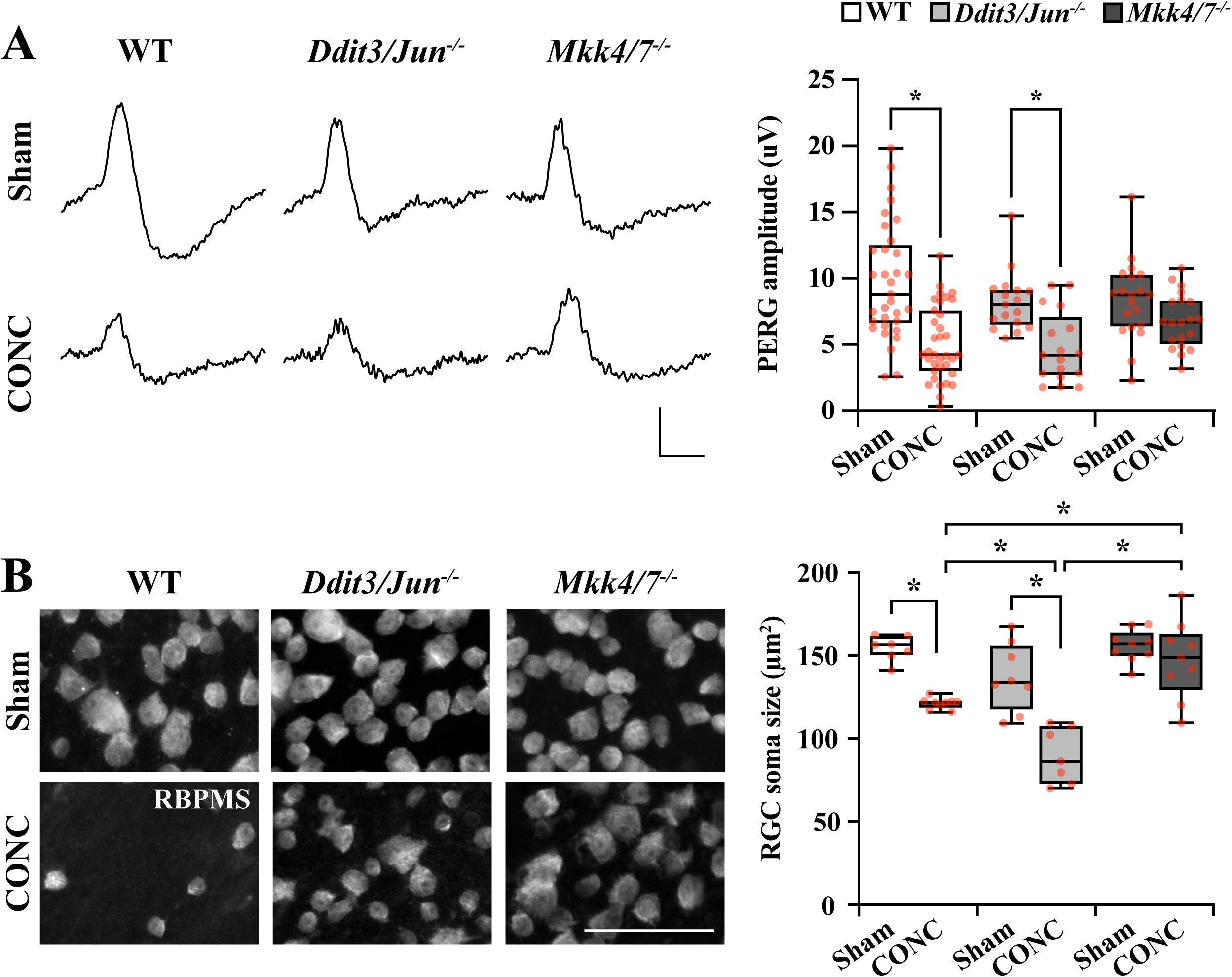
| *Mkk4/7* deletion, but not *Ddit3/Jun* deletion, prevented PERG amplitude decline and somal shrinkage after axonal injury. **A.** Representative PERG traces and quantification of PERG amplitudes from WT, *Ddit3/Jun^-/-^,* and *Mkk4/7^-/-^* retinas 14 days post-CONC or Sham procedures. WT and *Ddit3/Jun^-/-^* PERG amplitudes significantly declined 14 days post-CONC (by 52.6±5.7, \**P*<0.001 and 57.9±7.9µV, \**P*=0.002, respectively), while *Mkk4/7^-/-^* PERG amplitudes did not significantly decline (*n≥*17, two-way ANOVA, Holm-Sidak’s *post-hoc*.). Scale bar: Y: 5μV, X: 100ms. **B.** Representative high-resolution images of retinal flat mounts immunoassayed for RBPMS (scale bar, 50μm) and quantification of RGC soma size from WT, *Ddit3/Jun*^-/-^ and *Mkk4/7^-/-^* retinas 14 days post-Sham or CONC procedures. After CONC, WT and *Ddit3/Jun^-/-^* RGC soma sizes were reduced to 78.4±0.8% (\**P*=0.001) and 65.5±4.6% (\**P*<0.001) the size of respective RGCs after Sham procedures. *Mkk4/7^-/-^* RGC soma sizes did not change after CONC compared to Sham (*P*=0.586). *Mkk4/7^-/-^* RGCs were significantly larger compared to WT (\**P*=0.006) and *Ddit3/Jun^-/-^* (\**P*<0.001) RGCs after CONC, and *Ddit3/Jun^-/-^* RGCs were significantly smaller than WT post-CONC (\**P*=0.002). Soma size (μm^2^)±SEM from WT, *Ddit3/Jun^-/-^,* and *Mkk4/7^-/-^* mice, respectively: Sham: 154.5±2.8, 137.0±7.3, 155.8±3.2; CONC: 121.2±1.1, 89.7±6.3, 147.5±7.8 (*n*≥7, two-way ANOVA, Holm-Sidak’s *post hoc*).

## Discussion

In glaucoma, injury to RGC axons drives degenerative signaling cascades that are critical for eliciting degeneration of the RGC soma and axon. The mechanisms governing RGC degeneration act in a compartment-specific manor (1, 2, 4, 29, 30). BAX-dependent apoptosis governed degeneration of the RGC proximal segment, but did not contribute to axonal degeneration (1). Several lines of evidence have suggested axonal degeneration mechanisms are critical drivers initiating cell death pathways ultimately driving degeneration of all RGC compartments (5, 7). Therefore, mechanisms important in axonal degeneration that can lead to downstream somal BAX activation have been recent targets of investigation.

The present study investigated the degenerative mechanisms by which RGCs die after glaucoma-relevant injury. Specifically, we examined the dual role of DDIT3 and JUN and their upstream regulators MKK4/7 in controlling RGC death after glaucoma-relevant injury. DDIT3/JUN controlled the majority of RGC somal death in ocular hypertensive DBA/2J mice but did not control axonal degeneration. Despite playing a critical role in RGC somal loss, DDIT3 and JUN did not contribute to decline in somal viability as measured by RGC somal shrinkage and loss of PERG amplitudes in DBA/2J mice or after CONC. In contrast, *Mkk4/7* deficiency significantly lessened not only somal apoptosis but also PERG amplitude decline, somal shrinkage, and the rate of axonal degeneration after glaucoma-relevant axonal injury. Together, these data suggest activation of MKK4 and MKK7 is a critical early event after glaucoma-relevant injury which activates pathways governing degeneration of RGC axons and somas. Future studies should assess the importance of MKK4/7 in driving RGC degeneration after chronic ocular hypertension. Furthermore, identifying the upstream activators of MKK4/7 in the context of axonal injury and elucidating the DDIT3/JUN-independent downstream effectors of MKK4/7 driving loss of somal viability and axonal degeneration will be important next lines of investigation.

Upstream regulators of MKK4 and MKK7 have previously been implicated in driving axonal and somal degenerative cascades after glaucoma relevant injury. Several studies have suggested the importance of Dual leucine kinase (DLK, MAP3K12), an upstream activator of MKK4 and MKK7, in initiating cell death pathways important in somal and axonal degeneration (30, 45, 46). Inhibition of DLK lessened Wallerian degeneration after axonal injury *in vitro* (15, 16, 46) and *in vivo* (16, 46). DLK inhibition modestly protected RGC somas and axons in a model of inducible ocular hypertension (47). However, deletion of *Dlk* did not phenocopy *Mkk4/7* deletion’s protection to RGC axons after CONC (30). Importantly, *Dlk* deletion was not sufficient to prevent axonal JNK activation after CONC (30), suggesting *Dlk* is not the sole activator of MKK4 and MKK7 after mechanical optic nerve injury. For example, MKK4/7 are also known to be activated by the MAP3K LZK (48, 49). Thus, it is possible that multiple MAP3Ks contribute to MKK4/7 activation in the context of glaucoma-relevant injury. Regardless of their upstream activators, MKK4/7 together controlled death of somal and axonal compartments after axonal injury. Our findings suggest MKK4/7 drive somal loss via activation of DDIT3 and JUN. However, axonal degeneration, somal functional decline, and soma shrinkage must be driven by MKK4/7 via a mechanism independent of DDIT3/JUN activation.

Much research has begun to uncover the mechanisms important in governing Wallerian axonal degeneration—many of which may be controlled by MKK4/7 after glaucoma-relevant injury. For example, recent work has suggested the importance of JNK signaling in axonal degeneration. Loss of JNK1, 2, and 3 activation prevented axonal degeneration *in vitro* (16, 43, 46) and *in vivo* after axonal injury (46, 50), including after CONC (16), suggesting MKK4/7 mediate axonal degeneration via activation of the JNKs. Importantly, although JNK2/3 are the JNKs known to be expressed in the central nervous system, *Jnk2/3* deletion did not prevent axonal degeneration after CONC (30) or in DBA/2J mice (51). These data either indicate a critical role for JNK1 in mediating axonal degeneration or suggest MKK4/7 mediate axonal degeneration through mechanisms in addition to or independently of JNK activation.

MKK4/7 (potentially via JNK activation) may drive axonal degeneration by contributing to NMNAT2 depletion after axonal injury. Walker et al. showed *Mkk4/7* silencing prevented NMNAT2 degradation and subsequent axonal degeneration in cultured dorsal root ganglion cell after axotomy. Furthermore, this study showed both MKK4/7 activation and consequential NMNAT2 degradation acted upstream of pro-degenerative SARM1 activation to drive axonal degeneration (43) (NMNAT2 degradation has been shown to act upstream of SARM1 activation in several models of axon injury (15, 43, 52–55)). It has been suggested members of the MAPK cascade, including DLK, directly target NMNAT2 for degradation (14, 15). It is possible MKK4/7 and/or JNK1/2/3 directly target NMNAT2 for degradation, ultimately triggering SARM1 activation and allowing axonal degeneration after glaucoma-relevant injury. Alternatively, work done by Yang et al. suggested SARM1 activity and NMNAT2 depletion acts upstream of MKK4/7-JNK1/2/3 activation to drive Wallerian degeneration after axonal injury (16). However, recent work has shown *Sarm1* deletion does not phenocopy *Mkk4/7* deletion after CONC. While *Sarm1* deletion protected RGC axons, it did not prevent somal JUN activation and subsequent somal death (29). Thus, it remains more likely MKK4/7 drive axonal degeneration by facilitating NMNAT2 degradation and allowing pro-degenerative SARM1 activation. Future work should elucidate the mechanisms downstream of MKK4/7 activation that ultimately drive Wallerian degeneration in the context of glaucomatous injury.

An intriguing result of our study indicated glaucoma-relevant loss of RGC gross potentials and soma shrinkage are not merely consequences of somal loss. To date, few studies have investigated the compartment-specific RGC-intrinsic mechanisms by which RGCs undergo shrinkage and lose the ability to fire in response to light-evoked inner retinal neuronal signals. Interestingly, previous work suggested *Bax* deletion, despite providing protection against neuronal loss, did not prevent somal shrinkage after neurotrophic deprivation. In this study, soma shrinkage corresponded with indicators of metabolic stress such as loss of glucose uptake and decreased rate of protein synthesis—which were not prevented by *Bax* deletion (42). In contrast, inhibition of mixed lineage kinases (and downstream MKK4/7 and JNK signaling) preserved soma size and metabolic integrity (41). Consistent with our reported results, these data indicate MKK4/7 promote deterioration of somal health independently of JUN/DDIT3-induced BAX activation.

It is possible MKK4/7 mediate somal and axonal decline via DDIT3/JUN-independent NMNAT2 degradation. WLD^S^ and *Nmnat* gene therapy’s preservation of RGC electrical function in ocular hypertensive mice, suggesting a role for NMNAT2 in maintaining RGC somal viability (5, 7, 8, 56). It is also important to consider manipulations like WLD^S^ and *Nmnat1* gene therapy preserved both RGC somas and axons in DBA/2J mice (5–7, 57). This could suggest protection of both the RGC soma and axon is required for protection of somal viability after glaucoma-relevant injury. Work by Chou et al. showed temporarily blocking RGC retrograde axonal transport without injuring RGCs caused a significant and reversable reduction PERG amplitude (58). Thus, loss of axonal transport itself could lead to subsequent loss of somal gross potentials. This suggests preservation of axons (and therefore preservation of axon transport) by *Mkk4/7* deletion could also preserve RGC somal gross potentials.

MKK4/7 could also drive RGC degeneration via activation of DDIT3/JUN-independent mechanisms in the soma. For example, MKK4/7 may drive somal and axonal degeneration via activation of pro-apoptotic BH3 only proteins. JNK is known to phosphorylate and activate pro-apoptotic BH3 only proteins, thereby inhibiting pro-survival Bcl-2 family proteins and allowing BAX activation (17). Overexpression of *Bcl2l1* (*Bclxl*) prevented axonal degeneration in DBA/2J mice (28) and reduced RGC soma loss but not axonal degeneration after axonal injury (27, 59). Notably, *Bclxl* overexpression did not prevent somal shrinkage or PERG amplitude decline after axonal injury, suggesting BCLXL loss alone is not the sole driver of RGC somal degeneration in the context of glaucoma. *Bcl2* overexpression also protected RGC somas (60–62) but not distal axons (61, 62) after CONC. Interestingly, *Bcl2* overexpression prevented PERG amplitude decline even up to 2 months post-CONC (38), suggesting a potential role of Bcl2 family proteins in maintaining loss of somal viability after injury. Therefore, BCL2 or other Bcl-2 family proteins may be critical in maintaining both somal survival, somal health, and axonal degeneration, and MKK4/7-mediated inhibition of Bcl-2 family proteins may contribute to RGC degeneration in glaucoma.

In conclusion, we demonstrate MKK4/7 controlled loss of the RGC soma after glaucoma-relevant injury—likely via somal DDIT3/JUN activation. MKK4/7 also controlled RGC axonal degeneration and decline in somal viability via DDIT3/JUN-independent mechanisms. These data suggest MKK4/7 activation may be an early inciting mechanism initiating degenerative cascades in glaucoma. The DDIT3/JUN independent downstream mechanisms by which MKK4/7 drive axonal degeneration cascades, and deterioration of somal health remain unidentified. Future work should elucidate the downstream effectors of MKK4/7 in each RGC compartment and should also identify upstream activators of MKK4/7 in the context of glaucoma to further identify neurotheraputic targets.

## List of abbreviations

ACSF: artificial cerebrospinal fluid
CAP: compound action potential
cCASP3: cleaved caspase 3
CONC: controlled optic nerve crush
ERG: electroretinography
IOP: intraocular pressure
noe: no or early
PERG: pattern electroretinography
RGC: retinal ganglion cell
SEM: standard error of the mean
sev: severe
WT: wild type

## Acknowledgments

The authors would like to thank Drs. C. Tournier and R.J. Davis for generously providing the *Mkk4* and *Mkk7* floxed alleles. Thank you to Alyssa Parker and Thurma McDaniel for excellent technical assistance.

